# Warnings/Cautions when collecting Brassica diversity along a large climatic gradient

**DOI:** 10.1101/2023.09.15.557905

**Authors:** Cyril Falentin, Houria Hadj-Arab, Fella Aissiou, Claudia Bartoli, Giuseppe Bazan, Matéo Boudet, Lydia Bousset-Vaslin, Marwa Chwikhi, Olivier Coriton, Gwenaëlle Deniot, Julie Ferreira de Carvalho, Laurène Gay, Anna Geraci, Pascal Glory, Virginie Huteau, Riadh Ilahy, Vincenzo Ilardi, José A. Jarillo, Vladimir Meglič, Elisabetta Oddo, Mónica Pernas, Manuel Piñeiro, Barbara Pipan, Thouraya Rhim, Vincent Richer, Fulvia Rizza, Joëlle Ronfort, Mathieu Rousseau-Gueutin, Rosario Schicchi, Lovro Sinkovič, Maryse Taburel, Valeria Terzi, Sylvain Théréné, Mathieu Tiret, Imen Tlili, Franz Werner Badeck, Anne-Marie Chèvre

## Abstract

Agriculture faces great challenges to overcome global warming and to improve system sustainability, requiring access to novel genetic diversity. So far, wild populations and local landraces remain poorly explored. This is notably the case for the two diploid species, *Brassica oleracea* L. (CC, 2n=2x=18) and *B. rapa* L. (AA, 2n=2x=20). In order to explore genetic diversity in both species, we have collected numerous populations in their center of origin, the Mediterranean basin, on a large contrasting climatic and soil gradient from northern Europe to southern sub-Saharan regions. In these areas, we also collected 14 populations belonging to five *B. oleracea* closely related species. Before further genetic and agronomic investigations, we controlled the absence of species misidentification using flow cytometry, sequencing of species specific chloroplast genomic region, as well as cytogenetic analyses in case of unexpected results. Looking at the 102 *B. oleracea* and 146 *B. rapa* populations showing a good germination among the 112 and 154 populations collected, seventeen populations were misidentified. The most frequent mistake was a confusion of these diploid species with *B. napus*. Additionally for *B. rapa*, 2 autotetraploid populations were observed. Habitats of the collected wild populations and landraces are described in our work. This provides a unique plant material characterization that will pave the way for further analyses investigating the genomic regions involved in climatic and microbiota adaptation. This research is supported by the H2020 Prima project ‘BrasExplor’.

## Introduction

Agriculture has to face great challenges to overcome the global climate change and improve the sustainability of agricultural systems while maintaining crop production and quality. Regarding crop improvement, there are at least two main questions to consider: (*i*) which type of genetic diversity should we promote in breeding programs to withstand the new climatic regime and (*ii*) which material to select for the development of new relevant varieties in this erratic context. Intensive farming systems and particularly modern breeding techniques have led to a drastic reduction of the crop genetic diversity. On the other hand, local landraces and wild plant populations are a great source of genetic diversity. However, for many crop species such plant material has either never been collected, is not available, or has been poorly analysed and/or characterised.

The two diploid species that we focused on in this study, *Brassica oleracea* L. (CC, 2n=2x=18) and *B. rapa* L. (AA, 2n=2x=20), are particularly relevant model species for the analysis of diversity in relation with adaptation to the climate. They are native of the Mediterranean basin (Qi *et al*. 2017; Cheng *et al*. 2016; Bird *et al*. 2017; Cai *et al*. 2021; Mabry *et al*. 2021; McAlvay *et al*. 2021; Cai *et al*. 2022), in which they grow as wild populations or as local landraces selected over several generations by farmers. They encounter a large gradient of contrasted climate, soils and biotic factors from northern Europe to southern sub-Saharan regions. Exploring these wild populations and local varieties represents a unique opportunity to identify locally adapted material whose genetic diversity and adaptive traits could be relevant to face upcoming climate changes and disease emergences correlated to global change in the Mediterranean area, thus contributing to biodiversity-based agriculture.

Convergent evolution has lead to similar morphotypes in these two economically important vegetable species that were locally selected for a long time by farmers all over the Mediterranean basin, mainly for their heads (cauliflower or broccoli for *B. oleracea*, broccoletto for *B. rapa*), leaves (cabbage, kale for *B. oleracea;* fodder turnip for *B. rapa*) or roots (kohlrabi for *B. oleracea*, turnip for *B. rapa*). This morphological convergence between the two species is linked to their recent common ancestor (Cheng *et al*. 2016) as they diverged only 2-4My ago (Cheng *et al*. 2014). The morphological similarity between them is one of the reasons of some confusion when identifying the species. Additionally, a third species widely cultivated for seeds, resulting from the hybridization and genome doubling of the two diploid species, *B. napus* L. (AACC, 2n=4x=38), can also produce edible roots in swede cultivars, or leaves as fodder. As both species share many morphological characteristics with *B. napus*, species identification remains difficult and controls are required before further analyses.

In this paper, we describe the collection along a large climatic gradient of more than 100 populations of each *B. oleracea* and *B. rapa* species, including both landraces and wild populations. To ensure the absence of species misidentification or potential interspecific hybrids, plants of each population were assessed using different methods: (1) flow cytometry based on different genome size and chromosome number (630Mb for 18 chromosomes in *B. oleracea*, 529Mb for 20 chromosomes in *B. rapa*) (Belser *et al*. 2018), (2) Sanger sequencing of a species specific chloroplast genomic region (Li *et al*. 2017), and (3) cytogenetic approaches in the event of unexpected results from the previous analyses. After these controls, the geographical distribution and ecological environment of each population were described. This unique plant material will support further analyses from our consortium investigating the genomic regions involved in climatic and microbiota local adaptation.

## Material and methods

### Plant material

Wild populations of both *B. oleracea* and *B. rapa* species were collected in France. In addition, *B. rapa* wild populations were gathered in Italy, Algeria, Slovenia and *B. oleracea* in Spain. Pods were collected from 30 plants per population (when available), depending on the size and accessibility of the populations. Some wild populations of *B. oleracea* closely-related species were identified and added to the analysis: eight *B. montana* Pourr. populations (six from France and two from Italy), as well as two *B. rupestris* Raf. (subsp. *rupestris*), two *B. villosa* Biv. [subsp. *drepanensis* (Caruel) Raimondo & Mazzola and subsp. *tineoi* (Lojac.) Raimondo & Mazzola], one *B. macrocarpa* Guss., and one *B. incana* Ten., all from Sicily, Italy. *B. oleracea and B. rapa* landraces were collected in five different countries either through direct collects in farms for Algeria, Tunisia, Italy or in Biological Resource Centers maintaining old landraces in France (BRC BrACySol) and Slovenia (BRC KIS). In agreement with each country policy, the Nagoya treaty will be applied for the availability of the material.

Each collected population was named following a specific code. It starts with (i) two letters representing the species (BO for *B. oleracea*, BR for *B. rapa*, BM for *B. montana*, BU for *B. rupestris*, BV for *B. villosa*, BA for *B. macrocarpa*, and BI for *B. incana*), followed (ii) by a letter for the country of origin (F for France, I for Italy, S for Slovenia, E for Spain, A for Algeria, or T for Tunisia), (iii) then four letters indicating the location of the collecting site, (iv) either a W for a wild population or a L for a landrace, (v) and an additional letter (A, B, C, etc.) in case of several collecting sites at the same location (i.e. BR_I_CAST_W_A and BR_I_CAST_W_B). For all these populations, a common sheet was filled for wild populations to describe the environment (Suppl Table 1) and another one for landrace collects at the farm or when seeds were acquired from Genetic Resource Centers (GRC BrACySol in France, KIS in Slovenia). These data were synthesized in separate tables (Suppl Table 2).

**Table 1:** Origin and number of collected *B. rapa* and *B. oleracea* populations (including some closely related species). The number of populations, for which we obtained a germination sufficient for its multiplication, is indicated. For these latter, we also show the number of populations for which the species was validated using flow cytometry, chloroplast sequencing, plus cytogenetic controls when required.

| ***Brassica oleracea* and relatives species** |  |  |  |  |
| --- | --- | --- | --- | --- |
| Wild type |  |  |  |  |
| Species | Country | collected populations | Populations with a satisfying germination | validated populations |
| *Brassica oleracea* | France | 34 | 34 | 33 |
|  | Spain | 11 | 11 | 11 |
| *Brassica rupestris* | Italy | 2 | 2 | 2 |
| *Brassica incana* | Italy | 1 | 1 | 1 |
| *Brassica montana* | France | 6 | 6 | 6 |
|  | Italy | 2 | 2 | 2 |
| *Brassica macrocarpa* | Italy | 1 | 1 | 1 |
| *Brassica villosa* | Italy | 2 | 2 | 2 |
| Landrace |  |  |  |  |
| Species | Country | collected populations | Populations with a satisfying germination | validated populations |
| *Brassica oleracea* | Algeria | 11 | 7 | 7 |
|  | France | 29 | 29 | 28 |
|  | Italy | 6 | 5 | 5 |
|  | Slovenia | 17 | 13 | 13 |
|  | Tunisia | 4 | 3 | 3 |
| ***Brassica rapa*** |  |  |  |  |
| Wildtype |  |  |  |  |
| Species | Country | collected populations | Populations with a satisfying germination | validated populations |
| *Brassica rapa* | Algeria | 29 | 28 | 28 |
|  | France | 22 | 22 | 17 |
|  | Italy | 20 | 17 | 17 |
|  | Slovenia | 4 | 4 | 1 |
|  | Tunisia | 2 | 2 | 0 |
| Landrace |  |  |  |  |
| Species | Country | collected populations | Populations with a satisfying germination | validated populations |
| *Brassica rapa* | Algeria | 33 | 31 | 29 |
|  | France | 30 | 30 | 27 |
|  | Italy | 6 | 5 | 5 |
|  | Slovenia | 2 | 2 | 2 |
|  | Tunisia | 6 | 5 | 5 |

Thirty plants per population were grown in the greenhouse. For wild populations, we planted one seed of each of the 30 collected mother plants. When seeds were collected from less than 30 plants could be collected, we sowed seeds to equally represent each mother plant. For landraces, 30 seeds were sown.

As controls for the different experiments, we used a known representative of *B. oleracea, B. rapa* and *B. napus* species: doubled haploid lines of *B. oleracea* subsp. *italica* (HDEM) and *B. rapa* subsp. *trilocularis* (Z1) (Belser *et al*. 2018) and a pure line of *B. napus* subsp. *oleifera*, Darmor.

### Cytogenetic control and chromosome counts

Flow cytometry was performed on all plants for assessing the chromosome number of each plant using leaves as described by Leflon *et al*. (2006).

For some populations, the chromosome number was also determined from mitotic chromosomes observed on metaphasic cells isolated from root tips. Roots tips of 0.5 - 1.5 cm in length were treated in the dark with 0.04% 8-hydroxiquinoline for 2h at 4°C followed by 2h at room temperature to accumulate metaphases. They were then fixed in 3:1 ethanol-glacial acetic acid for 48h at 4°C and stored in ethanol 70 % at −20 °C until use. After being washed in distilled water for 10 min, HCl 0.25 N 10 min, then treated 15 min with a 0.01 M citric acid-sodium citrate pH 4.5 buffer, root tips were incubated at 37°C for 30 min in a enzymatic mixture (5% Onozuka R-10 cellulase (Sigma), 1% Y23 pectolyase (Sigma)). The enzymatic solution was removed and the digested root tips were then carefully washed with distilled water for 30 min. One root tip was transferred to a slide and macerated with a drop of 3:1 fixation solution. Dried slides were then stained by a drop of 4’,6-diamidino-2-phenylindole (DAPI). Cells were viewed with an ORCA-Flash4 (Hamamatsu, Japan) on Axio Imager Z.2 (Zeiss, Oberkochen, Germany) and analysed using Zen software (Carl Zeiss, Germany).

### Fluorescence *in situ* hybridization (FISH)

The BoB014O06 BAC clone from *B. oleracea* BAC library (Howell *et al*. 2008) was used as “genomic *in situ* hybridization (GISH)-like” to distinguish specifically all C-genome chromosomes in *B. napus* (Suay *et al*. 2014). The BoB014O06 clone was labelled by random priming with Alexa-594 dUTP. The ribosomal probe 45S rDNA used in this study was pTa 71 (Gerlach and Bedrook, 1979) which contained a 9-kb *Eco*RI fragment of rDNA repeat unit (18S-5.8S-26S genes and spacers) isolated from *Triticum aestivum* L. pTa 71 was labelled by random priming with biotin-14-dUTP (Invitrogen, Life Technologies). Biotinylated probes were immunodetected by Fluorescein avidin DN (Vector Laboratories). The chromosomes were mounted and counterstained in Vectashield (Vector Laboratories) containing 2.5µg/mL 4’,6-diamidino-2-phenylindole (DAPI). Fluorescence images were captured using an ORCA-Flash4 (Hamamatsu, Japon) on an Axio Imager Z.2 (Zeiss, Oberkochen, Germany) and analysed using Zen software (Carl Zeiss, Germany).

### Species identification using chloroplast sequences

The aim was to amplify a chloroplast genomic region containing diagnostic SNPs or indels for *B. oleracea, B. rapa* or *B. napus*. DNA of one to three plants per population and of control lines was extracted using 50mg of fresh leaf tissue, which had previously been freeze-dried, and the Nucleospin Plant II kit (Macherey Nagel). The primers used were trnK-rps16_F (5’ CATAAACAGGTAGACTGCTAACTGG 3’) and trnK-rps16_R (5’ GTATTCTTCCTAAAGGTATGAAAACTAAC 3’) with following PCR reagents: 1X buffer, MgCl2 2mM, dNTPs 0.25mM, Primers 0.5µM each, Taq Promega 1.5U and 5ng DNA of the sample analysed. The PCR conditions were a denaturation 94°C 2 min, then 35 cycles 94°C 30 sec - 59°C 30 sec - 72°C 1 min 30 sec with a final elongation 72°C 10 min. The amplified region was then sequenced by Sanger (Genoscreen) and analysed using Geneious software (https://www.geneious.com).

## Results

### 1. Analyses of the collected populations

Among the collected populations (Table 1), the first limiting factor encountered was the germination of the collected seeds, even under favorable controlled conditions applied on automated germination tools for *B. rapa*. Specifically, 6.8% of the collected populations showed a poor germination with less than 30 plants per population and were not considered for further analyses. This low germination may be attributed to two different factors: the high level of seed dormancy (observed here in 18.2% of *B. rapa* wild populations) and the seed conservation of landraces (5.2% and 14.9% of seeds showed a very poor germination rate for *B. rapa* and *B. oleracea* landraces, respectively).

To validate the correct species identification of each collected population and to verify the absence of contamination in the collected seeds, we performed flow cytometry on all the plants grown to represent a population. As the different investigated species have a profile linked to their differences in DNA content (630Mb for 18 chromosomes in *B. oleracea*, 529Mb for 20 chromosomes in *B. rapa*) (Figure 1A, B), it was possible to determine with +/-2 chromosomes the genomic structure of each plant.

**Figure 1.**
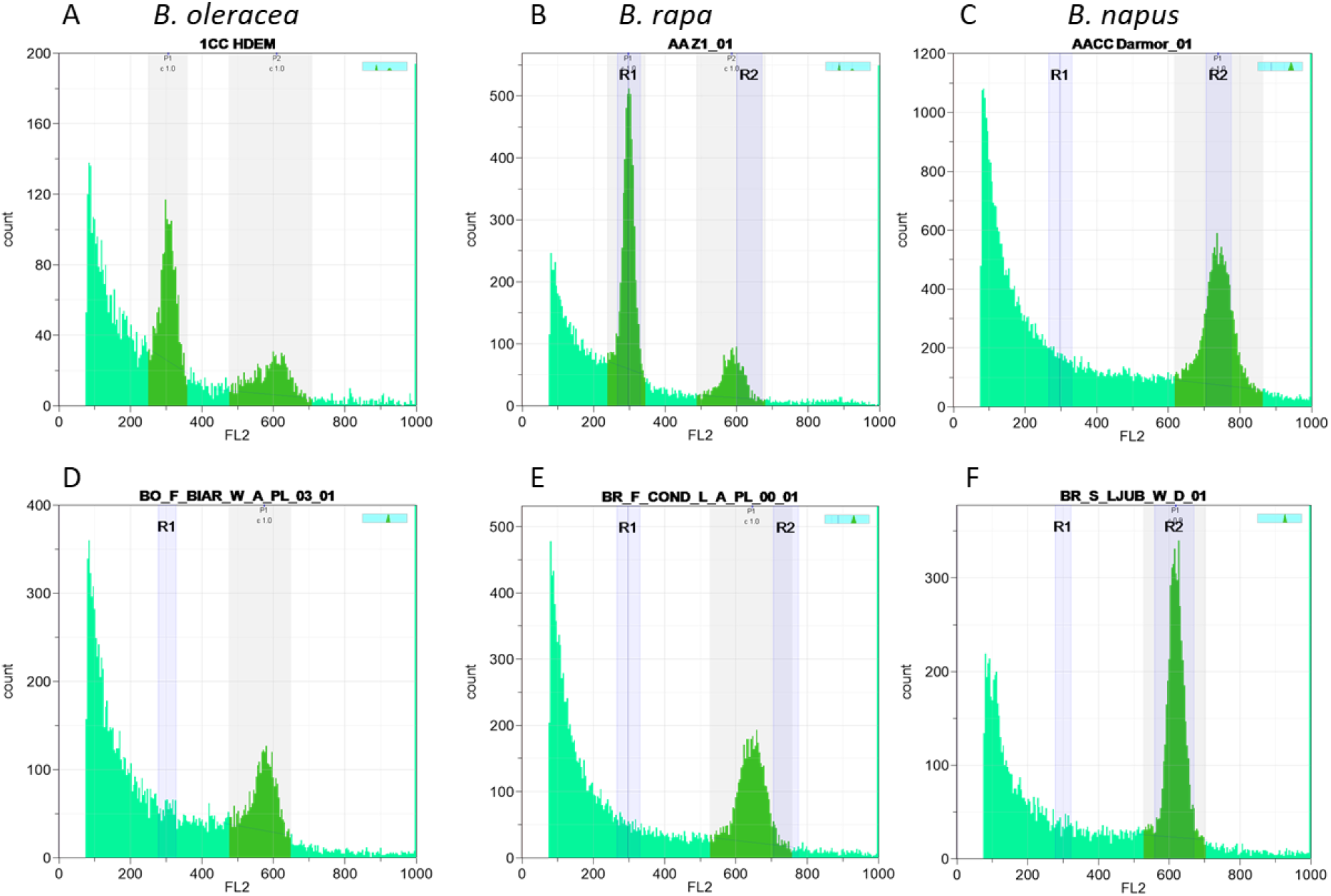
Flow cytometry identification of few populations harboring an unexpected profile compared to control populations: A) *B. oleracea*; B) *B. rapa*; C) *B. napus*; D) a misidentified *B. oleracea* population corresponding to *B. napus*; E and F) *B. rapa* autotetraploid populations.

However, due to possible contamination with species having a close chromosome number, this analysis was complemented by sequencing a chloroplast genomic region that showed species specific differences. The size of the amplified regions was 1118bp, 1088bp and 1084bp for *B. oleracea, B. rapa* and *B. napus*, respectively. After aligning the sequences, we compared the sequences with the controls. We observed four SNPs and six indels specific of *B. oleracea*, four SNPs and four Indels specific of *B. rapa* and three SNPs and five Indels specific of *B. napus* (examples provided in Figure 2A, B). *B. montana* (2n=18) differs from *B. oleracea* at only three SNPs and one Indel whereas *B. villosa* and *B. macrocarpa* differ from *B. oleracea* at seventeen SNPs and six Indels. *B. rupestris* showed exactly the same sequence as these two later species except for one SNP, indicating that these three species (*B. villosa, B. macrocarpa and B. rupestris*) are highly related to each other whereas *B. montana* seems closer to *B. oleracea* (Figure 2C).

**Figure 2.**
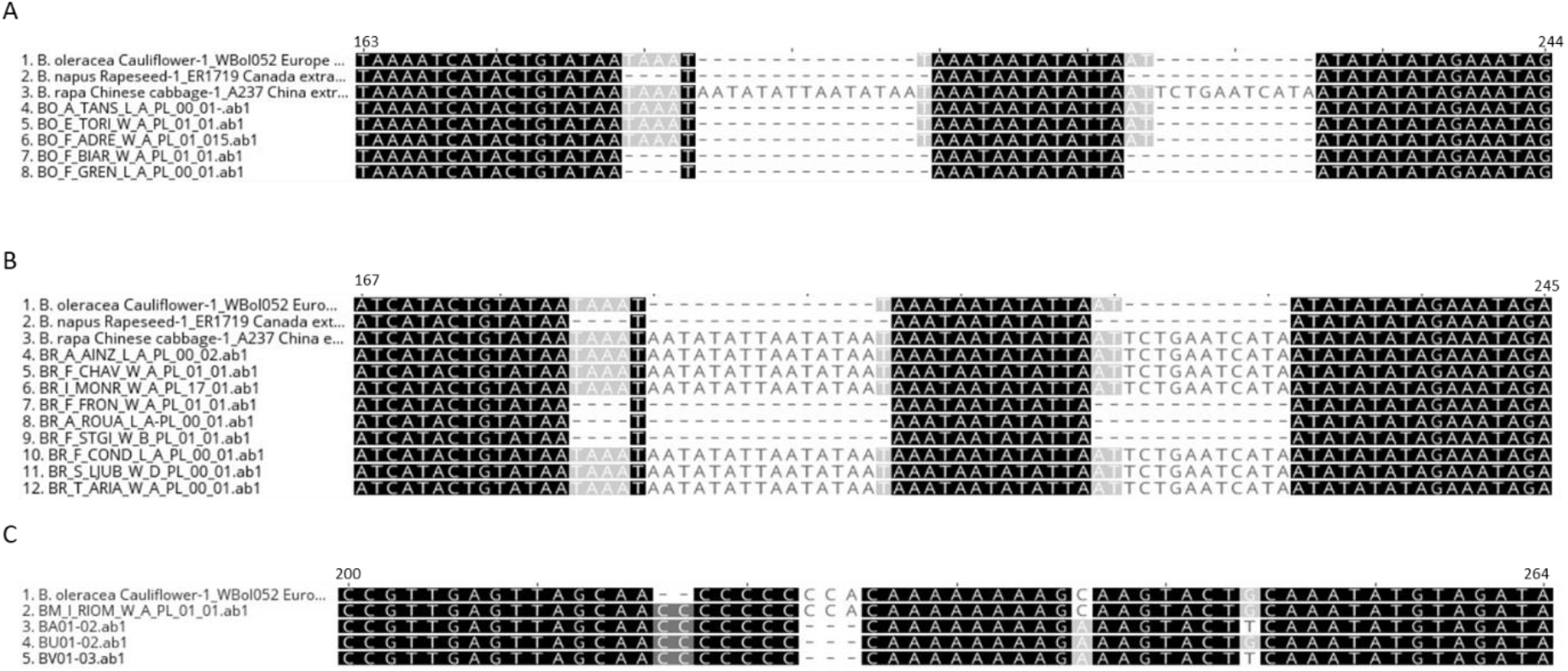
Examples of chloroplast regions showing differences between the species: (A) comparison between the controls and different *B. oleracea* populations, (B) comparison between the controls and different *B. rapa* populations, (C) comparison between the controls and different *B. oleracea* related species, *B. montana* (BM), *B. macrocarpa* (BA), *B. rupestris* (BU) and *B. villosa* (BV).

When flow cytometer and sequencing data were not congruent, chromosome counting was performed during mitosis. This observation was combined with GISH-like allowing identification of the C chromosome and of rDNA locus number, specific of each species with four, ten and twelve loci for *B. oleracea, B. rapa* and *B. napus*, respectively (Figure 3).

**Figure 3.**
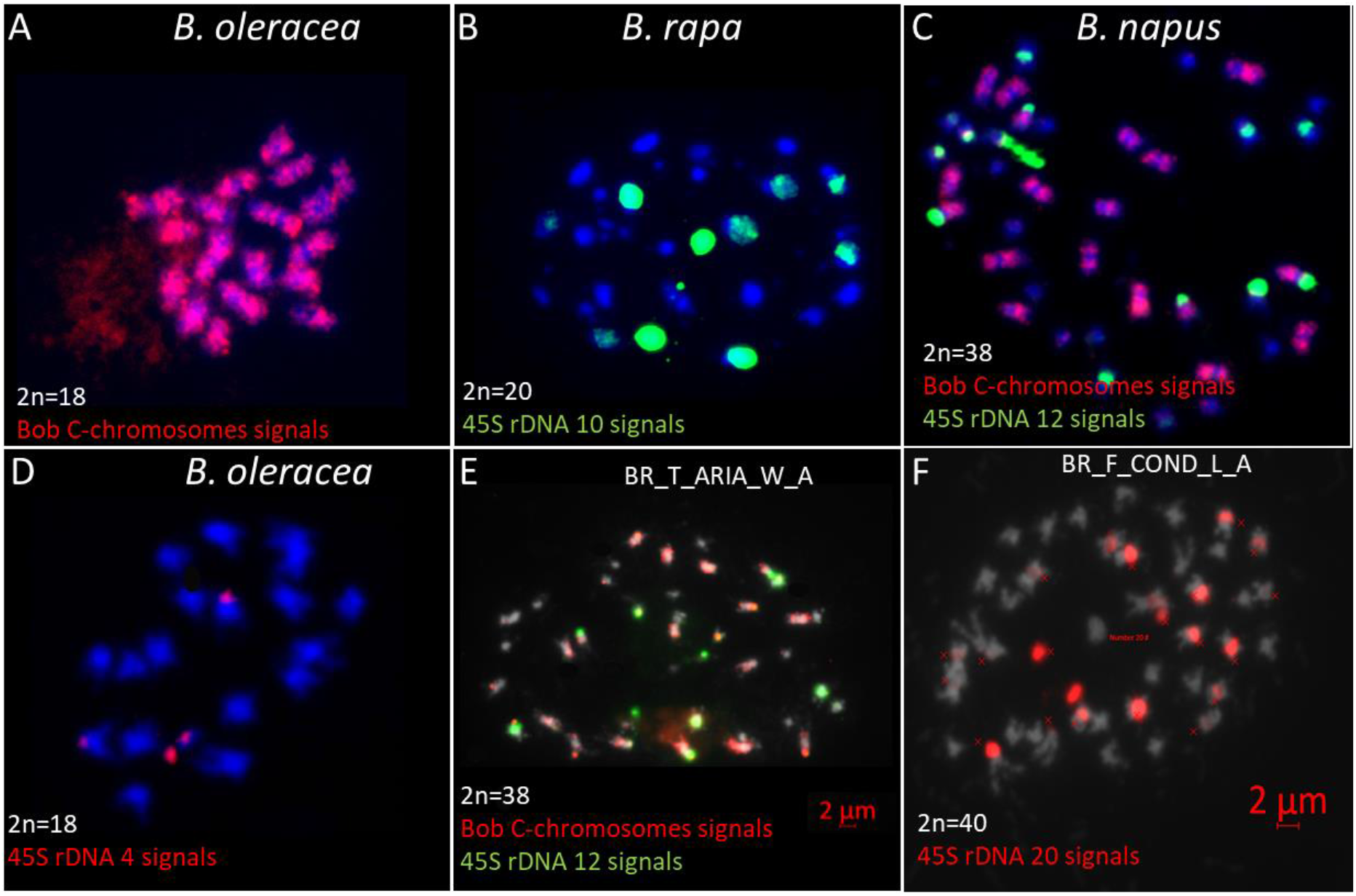
Chromosome number counted in mitosis with the three controls (*B. oleracea* A and D, *B. rapa* B and *B. napus* C) and two populations showing an unexpected structure: BR_T_ARIA_W_A (E) with *B. napus* genomic structure and BR_F_COND_L_A (F) an autotetraploid of *B. rapa*.

The most frequent mistake was a confusion between *B. oleracea* or *B. rapa* with *B. napus*. Among the 102 *B. oleracea* populations analysed, only two were misidentified (one wild and one landrace) and were thereafter confirmed to belong to *B. napus* using flow cytometry (Figure 1D for example). This misidentification was also validated by chloroplast sequencing (Figure 2A) and chromosome counting. Among the 146 analysed *B. rapa* populations, fifteen populations were misidentified, out of which twelve populations were identified as *B. napus*. Nine of these twelve populations were sampled in the wild and are probably volunteers of *B. napus*, i.e. escaped from the fields. All these data were confirmed by the sequencing of a chloroplast genomic region (Figure 2B) revealing that all carried *B. napus* chloroplasts except for one wild Tunisian population (BR_T_ARIA_W_A) which had *B. rapa* chloroplasts. The *B. napus* origin of this population was confirmed by cytogenetic analyses, revealing the presence of 9 C chromosomes and of 12 45S signals by FISH, 8 on A genome and 4 on C genome (Figure 3E). Among the three remaining misidentified *B. rapa* populations, one wild population from Tunisia had a cytometry value close to *B. rapa* but no chloroplast gene amplification was detected; further morphological observations of this population revealed that it probably belongs to the genus *Sinapis*. The two last cases observed were *B. rapa* populations (one Slovenian wild population BR_S_LJUB_W_D and one French landrace BR_F_COND_L_A) having a flow cytometry values close to the one of *B. napus* (Figure 1E and 1F) but a *B. rapa* chloroplast genomic sequence. Using cytogenetics, we detected no C chromosomes after a GISH-like experiment and 20 45S rDNA were counted, i.e. five rDNA loci per A genome (Figure 3F), which lead us to the conclusion that these populations were in fact *B. rapa* autopolyploids (AAAA, 2n=4x=40).

Most of the populations confirmed as belonging to a specific species had an identical chloroplast sequence. Nevertheless, we observed a few SNPs specific of some populations. In *B. oleracea*, two SNPs were specific to only seven populations (BO_F_JOUY_L_A, BO_S_LJUB_L_G, BO_S_LJUB_L_H, BO_S_LJUB_L_L, BO_S_LJUB_L_M, BO_S_LJUB_L_N and BO_S_LJUB_L_O) and one allele at a different SNP was specific to BO_F_MERS_W_A. In *B. rapa*, three variations differentiated a few populations, one SNP in BR_A_DELL_W_A, one base deletion in BR_A_SEBA_W_A and BR_A_BOME_W_A and one SNP in BR_A_BLID_W_A, BR_A_BOUF_W_A, BR_A_CHLE_W_A, BR_A_BARA_W_A. These differences were observed in all the individuals tested per population.

For *B. oleracea* related species (*B. montana, B. rupestris, B. villosa, B. macrocarpa* and *B. incana*), all collected populations per species had the same flow cytometry value and the same chloroplast sequence.

### 2. Description of the populations

After discarding the few populations that were misidentified, we further characterized the remaining populations and their respective data collected during harvest.

Wild *B. oleracea* populations were collected on cliffs on the atlantic coast in France and in Spain (Figure 4), whereas its related species (*B. montana, B. rupestris, B. villosa, B. macrocarpa* and *B. incana*) were growing more in Southern regions, on the mediterrannean coast. Their locations and the characteristics of each environment are described in Suppl. Table 1.

**Figure 4.**
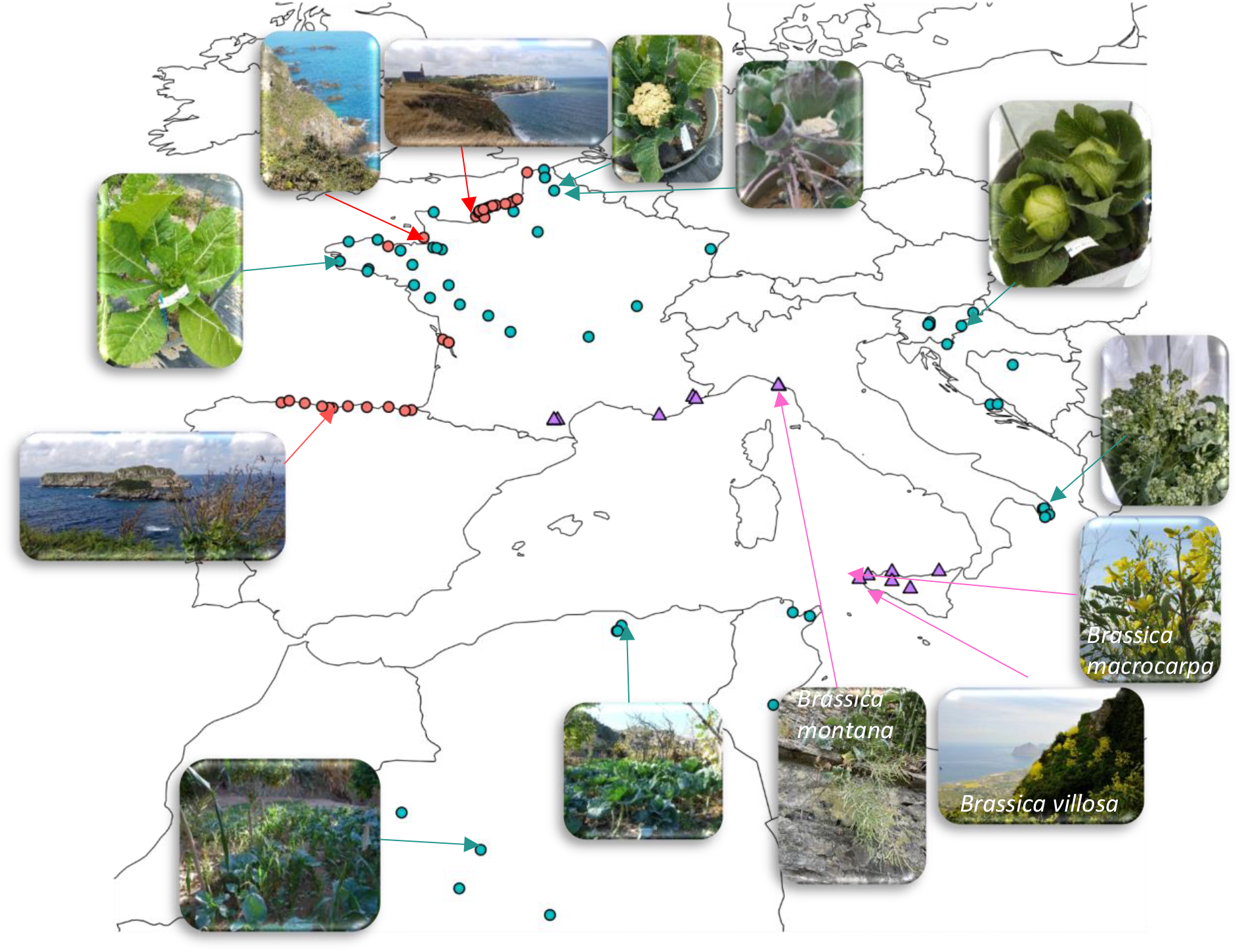
Distribution of the *B. oleracea* populations collected: 44 wild populations indicated with red dots, 56 landraces with green dots and 14 related species with pink triangles. The pictures illustrate the different plant morphology as well as the diversity of the original landscapes.

*B. oleracea* landraces were selected by farmers in each country, even in very warm regions such as the South of Algeria (Figure 4; Suppl Table 2). Selection of different organs for crop production (flowers, leaves, stems or roots) has lead to the divergence of highly diverse phenotypes. It is worth to mention that some morphotypes are difficult to classify in one subspecies as some of them were domesticated at the same time for leaf production as subsp. *acephala* and for head cabbage as subsp. *capitata* (i.e. BO_A_TAZL_L_A). Additionally, even within the same morphotype, different developmental traits can be observed such as in Mugnoli populations (south of Italy) with several floral heads compared to common broccoli (Laghetti *et al*.2005).

Wild *B. rapa* populations (Figure 5; Suppl Table 1) were found in locations where competition with other species is lower, such as vineyards, orchards or field margins. Thus, regardless of the country, the populations were generally large.

**Figure 5.**
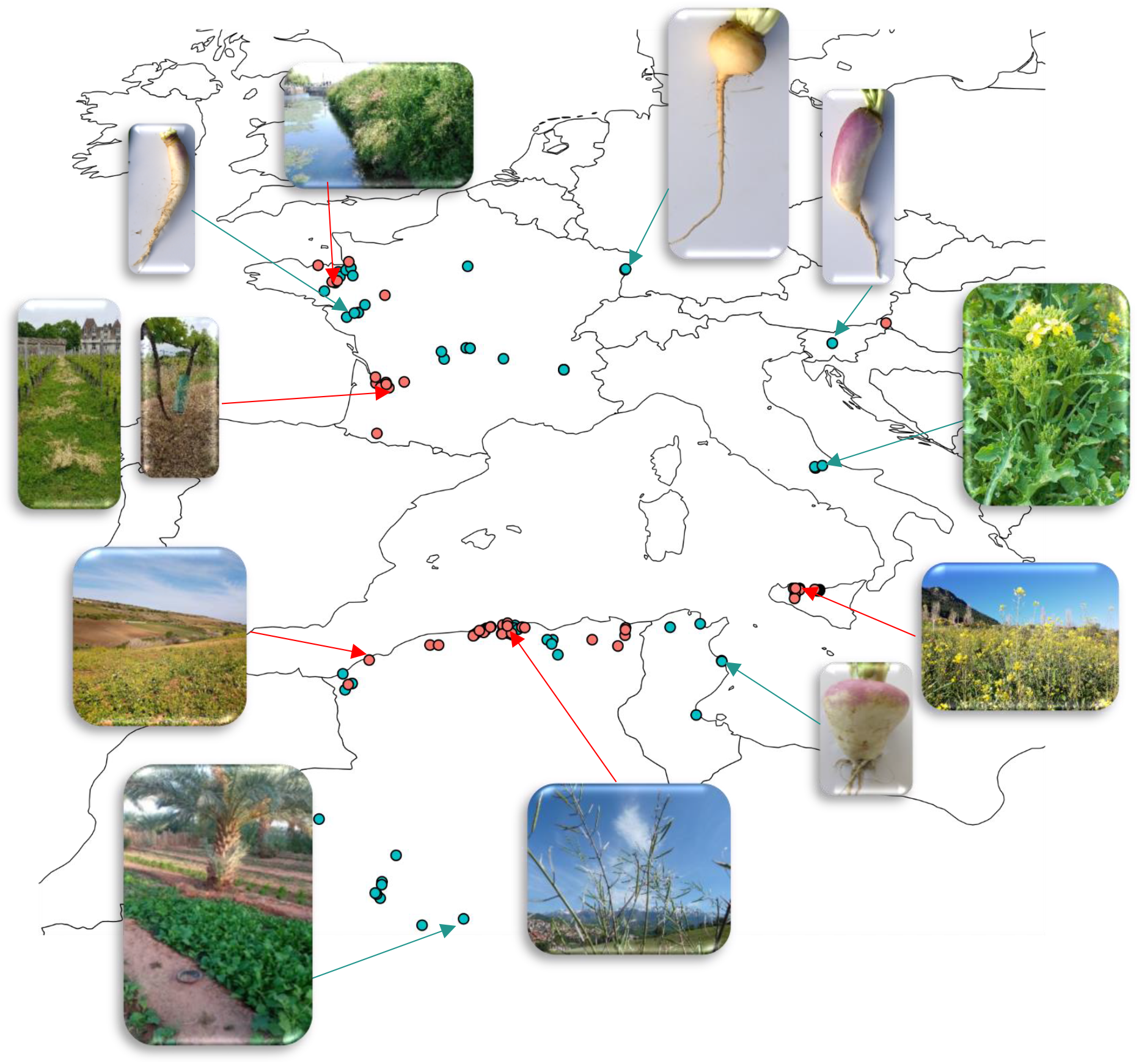
Distribution of the *B. rapa* populations collected: 63 wild populations indicated with red dots, 68 landraces with green dots. The pictures illustrate the different plant morphology as well as the diversity of the original landscapes.

The majority of the collected *B. rapa* local landraces (Figure 5; Suppl Table 2) were turnips (subsp. *rapa*) with the exception of few broccoletto (subsp. *sylvestris* var. *esculenta*) selected by Italian farmers.

## Discussion

In this paper, we described the sampling of wild populations and local landraces of *B. oleracea* and *B. rapa* along a large climatic and soil gradient from the North of France to the Sub-Saharan regions. Our objective was to validate that the seeds collected from plants of 112 and 154 populations (both wild and local landraces) of *B. oleracea* and *B. rapa*, respectively, belonged to the expected botanical species before beginning genetic studies of plant adaptation.

The first limiting factors was the germination. Seed dormancy was only detected among *B. rapa* populations. This trait, described in *Brassica* as a primary physiological dormancy (Finch-Savage and Leubner-Metzger, 2006), seems to be a characteristic of few wild *B. rapa* populations. The conditions of seed conservation on the other hand is a likely explanation for the low germination rate in landraces of both species. This observation highlights the importance of seed quality and storage conditions, especially in Genetic Resource Centers (Subramarian *et al*. 2023).

Because of the morphological similarity between the species, our controls have revealed the importance of performing molecular and cytogenetic analyses before undertaking genetic and agronomic studies. Flow cytometry is a high throughput technique allowing DNA content assessment of all plants, i.e. 30 plants per population. Yet, as several species of the *Brassiceae* tribe have a similar DNA content, this technique might not be precise enough (Leflon *et al*. 2006) to validate the species. That is the reason why we completed this analysis by sequencing a species-specific chloroplast genomic region taking advantage of the whole chloroplast genome sequences of many Brassica species/populations published by Li *et al*. (2017). The combination with the analysis of chloroplast sequences allowed the confirmation of a mistake for one Tunisian population presenting a flow cytometry value similar to *B. rapa* but no chloroplast amplification as it probably belongs to the genus *Sinapis*. However, the most frequent mistake was a confusion with *B. napus*, showing a higher DNA content, detectable by flow cytometry. Thus, among the fourteen populations determined as *B. napus* by flow cytometry (two populations in the *B. oleracea* and twelve in *B. rapa*), three had a chloroplast sequence similar to *B. rapa*. That is the explanation why cytogenetic experiments were performed on plants of these three populations, using GISH-like on mitotic chromosomes with a BAC specific of *B. oleracea* chromosomes (Suay *et al*. 2014) and 45S rDNA probes revealing the number of rDNA loci (Książczyk *et al*. 2011). From this data, we concluded that one Tunisian population was a *B. napus* population. It could be interesting to precisely compare after chloroplast assembly with the results reported by Li *et al*. (2017) who reported that *B. napus* chloroplasts can be classified into the two different clades identified from different *B. rapa* morphotypes. The two other populations were *B. rapa* autotetraploids, with 40 A chromosomes and 20 45S loci as expected when doubling the A genome. Such autopolyploid populations were previously reported for the production of new forage varieties (Olsson and Ellerström, 1980).

Among the 100 and 131 confirmed populations for *B*.*oleracea* and *B. rapa* respectively, chloroplast sequences revealed only a few variants (SNPs or Indels) for some accessions in both species. The low mutation rate of the chloroplast DNA in most flowering plant families can explain these variations as already reported from global chloroplast assembly. Interestingly, Li *et al*. (2017) observed more SNVs (Single Nucleotide Variants) in the *B. rapa* than in the *B. oleracea* genotypes that they investigated, with 343 and 16 SNV, respectively.

By investigating an enlarged *B. oleracea* diversity, Perumal *et al*. (2021) described more SNVs with clustering of different cultigroups. In our collected wild and landraces populations, we observed that a common variation is shared by seven populations belonging to *capitata* and *acephala* groups originating from Slovenia with the exception of one French landrace. For *B. rapa*, SNV were only observed in some wild Algerian populations. Further studies are in progress in order to compare the genetic diversity from chloroplast assembly and nuclear SNP polymorphism, taking into account the different cultigroups and their geographic origins.

A large morphological diversity was observed among the *B. oleracea* landraces whereas wild populations were morphologically similar to forage kales. For Mugnoli belonging to the same group as broccoli (subsp. *italica*), Biancolillo *et al*. (2023) developed a non-destructive tool based on Multivariate Image analysis and agro-morphological descriptors for the characterization and authentication of these local varieties. For *B. rapa*, landraces selected by farmers are mainly turnips, with the exception of five populations of Broccoletto. In this paper, we described the different environments in which these different populations were collected.

This well characterized material collected on a very large climatic and soil gradient opens the prospect to identify genomic regions involved in adaptation to climatic constraints and microbiota descriptors. To do so, seeds were produced at the same geographic location in order to avoid a maternal effect on seed quality. Thereafter, we will be able to perform high-throughput sequencing for bulks of 30 plants per population to capture the maximum diversity existing within the population. Mapping on the reference genome of each species and SNP calling will allow the description of genetic diversity and the design of nested core collections. Genome-wide association (GWAS) and genotype-environment association (GEA) analyses will be possible from the project consortium to identify genomic regions involved in climate adaptation. Functional analyses will be performed on the most contrasted populations to finely investigate their responses to cold and warm temperatures. Field experiences of core collections in five countries will allow the validation of favorable alleles under different environmental conditions. All the research is funded by H2020 Prima, BrasExplor (https://brasexplor.hub.inrae.fr/). All these data will be used (1) to promote local landraces, as several are endangered, and (2) to design crosses that could be relevant to produce pre-breeding populations, each adapted to the climatic evolution of each country.

## Acknowledgements

We thank the Genetic Resource Centers BrACySol (https://www6.rennes.inrae.fr/igepp_eng/About-IGEPP/Platforms/BrACySol) and the Agricultural institute of Slovenia (https://www.kis.si/en/) for providing seeds from different landraces. We thank Biogenouest (the western French network of technology core facilities in life sciences and the environment, supported by the Conseil Regional des Pays de la Loire) for the access to the molecular cytogenetics (https://www6.rennes.inrae.fr/igepp_eng/About-IGEPP/Platforms/Molecular-Cytogenetics-Platform-PCMV) and GenOuest bioinformatic platforms (https://www.genouest.org/). We thank Plant imaging platform PHENOTIC in Angers (INRAE-IRHS, Angers University, Institut Agro, GEVES, France) for experiments on seed germination. We would also like to thank all the staff that took care of our plant material (especially L. Charlon, J-P. Constantin and F. Letertre).

All the work presented is supported by H2020 Prima project no. 1425, BrasExplor “Wide exploration of genetic diversity in Brassica species for sustainable crop production” and by INRAE through TSARA Initiative (Transforming food systems and agriculture through a partnership research with Africa) promoting a specific French-Algerian collaboration.

## Contribution

CF, HH, AMC designed and managed all the experiments. CF, HH, FA, CB, GB, LB, MC, GD, JFC, LG, AG, RI, VI, JAJ, VM, EO, MP,MP, BP, TR, FR, JR, RS, LS, VT, ST, IT, FWB, AMC participated to the collects and the local description of the populations. VM, BP,VR, ST provided landraces and their description from BRC. MT performed flow cytometer analyses. GD and MRG designed chloroplast markers and performed experiments. OC and VH performed all molecular cytogenetic experiments. MB managed the database for population description. CF, HH, MRG and MT contributed to writing the manuscript, which was finalized by AMC.

## Supplemental data

- Supplemental Table 1: Description of *B. oleracea* and *B. rapa* wild populations
- Supplemental Table 2: Description of *B. oleracea* and *B. rapa* landraces

